# An evolutionary genomic approach reveals both conserved and species-specific genetic elements related to human disease in closely related *Aspergillus* fungi

**DOI:** 10.1101/2021.03.01.433394

**Authors:** Matthew E. Mead, Jacob L. Steenwyk, Lilian P. Silva, Patrícia A. de Castro, Nauman Saeed, Falk Hillmann, Gustavo H. Goldman, Antonis Rokas

**Affiliations:** Department of Biological Sciences, Vanderbilt University, Nashville, Tennessee, USA; Faculdade de Ciências Farmacêuticas de Ribeirão Preto, Universidade de São Paulo, Ribeirão Preto, Brazil; Junior Research Group Evolution of Microbial Interactions, Leibniz Institute for Natural Product Research and Infection Biology, Hans Knöll Institute (HKI), Jena, Germany

**Author notes:** Author for Correspondence: Antonis Rokas, Department of Biological Sciences, Vanderbilt University, Nashville, USA. MEM and JLS contributed equally to this work.

**Keywords:** evolution, *Aspergillus fumigatus*, evolutionary rate, virulence, pathogenicity, fungal disease, aspergillosis, convergent evolution

## Abstract

Aspergillosis is an important opportunistic human disease caused by filamentous fungi in the genus *Aspergillus*. Roughly 70% of infections are caused by *Aspergillus fumigatus*, with the rest stemming from approximately a dozen other *Aspergillus* species. Several of these pathogens are closely related to *A. fumigatus* and belong in the same taxonomic section, section *Fumigati*. Pathogenic species are frequently most closely related to non-pathogenic ones, suggesting *Aspergillus* pathogenicity evolved multiple times independently. To understand the repeated evolution of *Aspergillus* pathogenicity, we performed comparative genomic analyses on 18 strains from 13 species, including 8 species in section *Fumigati*, which aimed to identify genes, both ones previously connected to virulence as well as ones never before implicated, whose evolution differs between pathogens and non-pathogens. We found that most genes were present in all species, including approximately half of those previously connected to virulence, but a few genes were section- or species-specific. Evolutionary rate analyses identified hundreds of genes in pathogens that were faster-evolving than their orthologs in non-pathogens. For example, over 25% of all single-copy genes examined in *A. fumigatus* were faster-evolving. Functional testing of deletion mutants of 17 transcription factor-encoding genes whose evolution differed between pathogens and non-pathogens identified eight genes that affect either fungal survival in a model of phagocytic killing, host survival in an animal model of fungal disease, or both. These results suggest that the evolution of pathogenicity in *Aspergillus* involved both conserved and species-specific genetic elements, illustrating how an evolutionary genomic approach informs the study of fungal disease.

## Introduction

The ability of a microbe to cause disease is a multi-factorial trait that is dependent upon a diverse set of genomic loci. For opportunistic fungal pathogens, whose “accidental” infections of humans are not a part of their normal life cycle (Casadevall and Pirofski 2007), the evolution of genomic loci contributing to virulence is thought to have been shaped by diverse evolutionary pressures, such as avoiding predation from soil-dwelling amoebae and surviving in warm and stressful environmental niches similar to those found inside human hosts (Tekaia and Latgé 2005; Nielsen et al. 2007; Hillmann et al. 2015). However, the evolution of fungal genes in the context of the evolution of fungal pathogenicity has yet to be fully studied (Sharpton et al. 2009; Gabaldón et al. 2016; Gupta et al. 2020; Rokas et al. 2020). This is especially true for filamentous fungi in the genus *Aspergillus*, which infect hundreds of thousands of human each year (Brown et al. 2012).

Aspergillosis, the spectrum of diseases caused by fungi in the genus *Aspergillus*, is a major global health issue and primarily afflicts individuals with compromised immune systems or who have other lung diseases or conditions (Gregg and Kauffman 2015). Approximately 70% of aspergillosis patients are infected with *Aspergillus fumigatus*, but a small number of infections are caused by other members of the genus (Alastruey-Izquierdo et al. 2014; Perlin et al. 2017; Latgé and Chamilos 2019). Some of these pathogenic species are very closely related to *A. fumigatus* and belong to the same taxonomic section, section *Fumigati* (Balajee et al. 2009; Alastruey-Izquierdo et al. 2013; Houbraken et al. 2020). In contrast, most of the approximately 60 species in section *Fumigati* do not cause disease or have rarely been found in the clinic, suggesting that the ability to cause disease evolved multiple times independently (or convergently) within *Aspergillus* (Rokas et al. 2020). For example, *Aspergillus oerlinghausenensis* and *Aspergillus fischeri*, the two closest relatives of *A. fumigatus* are both considered non-pathogenic (Houbraken et al. 2016; Mead et al. 2019; Steenwyk, Mead, Knowles, et al. 2020).

Why some *Aspergillus* species routinely infect humans whereas their very close relatives never or rarely do remains an open question (Rokas et al. 2020). To date, studies addressing this question have focused on comparing *A. fumigatus* to one or a few select species (Fedorova et al. 2008; Wiedner et al. 2013; Mead et al. 2019; Knowles et al. 2020; Steenwyk, Mead, Knowles, et al. 2020). Many individual genes and pathways are known to contribute to *A. fumigatus* virulence (Abad et al. 2010; Bignell et al. 2016; Brown and Goldman 2016; Steenwyk, Mead, de Castro, et al. 2020), but if they are present or function in the same manner in other section *Fumigati* species, including in non-pathogens, has rarely been studied (Knowles et al. 2020; Steenwyk, Mead, Knowles, et al. 2020). Genes associated with pathogenicity could be shared or absent amongst all pathogens, including *A. fumigatus*, following a “conserved pathogenicity” model, or be uniquely present (or absent) in each pathogen (“species-specific pathogenicity” model) (Rokas et al. 2020). The two models are not mutually exclusive, and the limited evidence available suggests that some genetic determinants of virulence likely follow the conserved pathogenicity model (Fedorova et al. 2008; Kjærbølling et al. 2018), whereas others follow the species-specific pathogenicity model (Fedorova et al. 2008; Kowalski et al. 2019; Mead et al. 2019).

To study the signatures of the repeated evolution of pathogenicity on those genes previously connected to *A. fumigatus* virulence as well as to all genes in *Aspergillus* genomes, we analyzed the genomes of 18 *Aspergillus* strains representing 13 species, including both pathogenic and non-pathogenic species from section *Fumigati*. We identified gene families across our entire genomic dataset and analyzed their presence/absence, family number distribution, and evolutionary rate in both pathogens and non-pathogens. After identifying multiple gene families with virulence-related genomic traits, we tested their *A. fumigatus* null mutants in multiple virulence assays and found both unique and previously identified factors as being important for pathogenicity-relevant traits. We found dozens of *A. fumigatus*-specific gene families but no pathogen-specific genes families, providing support for the species-specific pathogenicity model. However, we identified over 1,700 gene families that showed pathogen-specific evolutionary rates and fewer than 40 gene families with *A. fumigatus*-specific evolutionary rates, thereby supporting the conserved pathogenicity model. In addition, functional assays of deletion mutants of genes supporting either pathogenicity model identified eight genes that significantly affected pathogenicity-related traits. Our results provide support for both the conserved and species-specific pathogenicity models and show that previously identified virulence-related genes are largely conserved throughout section *Fumigati* and outgroups. More broadly, our study shows that an evolutionary genomic approach is a useful framework for gaining insights into the molecular mechanisms by which *Aspergillus* species impact human health.

## Results

### A genome-scale phylogeny of *Aspergillus* section *Fumigati*

Phylogenetic relationships among taxa in section *Fumigati* (Table S1) were examined using three maximum likelihood approaches – concatenation without gene-based partitioning, concatenation with gene-based partitioning, and coalescence – using 3,601 single copy orthologous genes. Both concatenation approaches yielded the same topology, recovering *A. neoellipticus* nested within *A. fumigatus* (Figure 1). All bipartitions received full support except the split between *A. neoellipticus* and *A. fumigatus* strains F16311 and 12-750544, which received 98% ultrafast bootstrap approximation support. The coalescence approach inferred a fully supported alternative topology that placed *A. neoellipticus* sister to the four *A. fumigatus* strains. Whether *A. neoellipticus* is conspecific with *A. fumigatus* or a distinct species has been previously discussed in the literature (Li et al. 2014) and our genome-scale analyses reflect this debate. Given the close evolutionary relationship of the two species, we choose to refer to *A. neoellipticus* as a strain of *A. fumigatus* rather than a distinct species.

**Figure 1.**
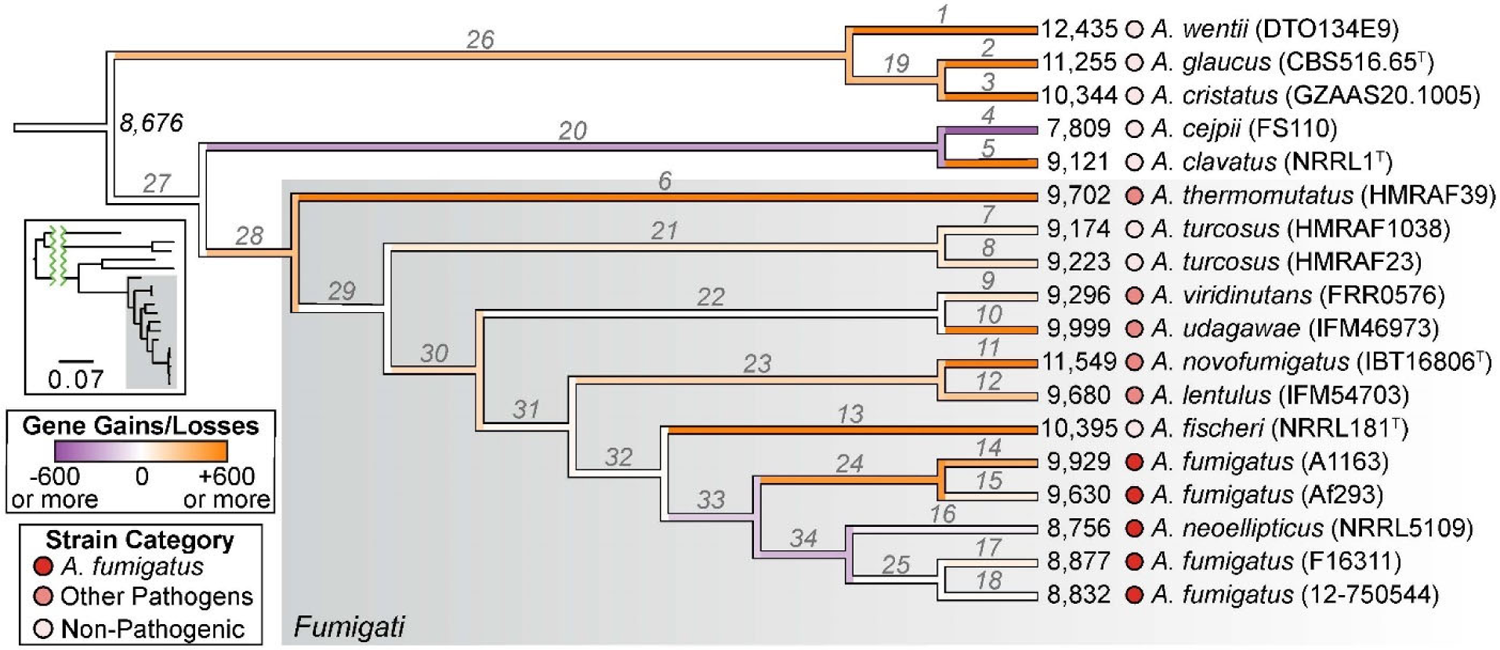
Genome-scale phylogeny and evolution of net gene gains or losses across *Aspergillus* section *Fumigati*. Relationships among taxa in section *Fumigati* inferred from a concatenation-based, maximum likelihood approach. Branches are colored based on the number of net gene gains or losses, and 8,676 genes were inferred at the last common ancestor of all taxa studied. Numbers at branch tips represent the total number of genes in that genome. The group of each strain is shown via shades of red (dark – most pathogenic, light – least pathogenic). Strain designations are in parenthesis next to species names and type strains are denoted by a superscript “T” next to their strain designations. Insert shows the phylogeny with branch lengths reflective of the estimated number of nucleotide substitutions per site (scale bar is 0.07 substitutions/site); taxa are in the same order as the larger cladogram. The number of gene gains, losses, and the net gain or loss are shown in Table S2 and use the branch labels shown here in light gray italics.

### Broad conservation of genes, including of virulence-related ones, across section *Fumigati*

To understand variation in the distribution of genes, including ones known to be involved in *A. fumigatus* virulence, we inferred gene and gene family gains and losses for every branch on the phylogeny (Figures 1 and S1). We inferred a net gain of 305 genes in the last common ancestor of section *Fumigati*. In addition, we found a net loss of 171 genes in the last common ancestor of *A. fumigatus* strains (Figure 1 and Table S2). An estimated net gain of 494 genes occurred in the last common ancestor of the two *A. fumigatus* reference strains, A1163 and Af293. The same general patterns of genome expansion and contraction were observed when gene family gain and loss were estimated (Supplementary Figure S1).

To test the validity of the conserved and species-specific pathogenicity models of virulence, we searched the 18 *Aspergillus* genomes for pathogen- or *A. fumigatus*-specific genes using both candidate and unbiased approaches. The candidate approach consisted of determining the presence or absence pattern of 206 virulence-related genes (Steenwyk, Mead, de Castro, et al. 2020) that our gene family analysis placed into 189 gene families. The number of gene members in each family ranged from 259 (in the gene family containing the transporter *abcC* – <u>Afu1g14330</u> (Paul et al. 2013)) to six (in the gene family containing the terpene cyclase *fma-TC* – <u>Afu8g00520</u> – from the fumagillin biosynthetic gene cluster (Guruceaga et al. 2018)). The same number of family members was found in every genome for 84 / 189 (∼44%) of the virulence-related gene families, including for 81 families with one copy in each strain. Of those virulence-related gene families that differed in family member size across the 18 genomes, there were no virulence-related gene families with only members in pathogens, section *Fumigati* species, or *A. fumigatus* (Figure 2A).

**Figure 2.**
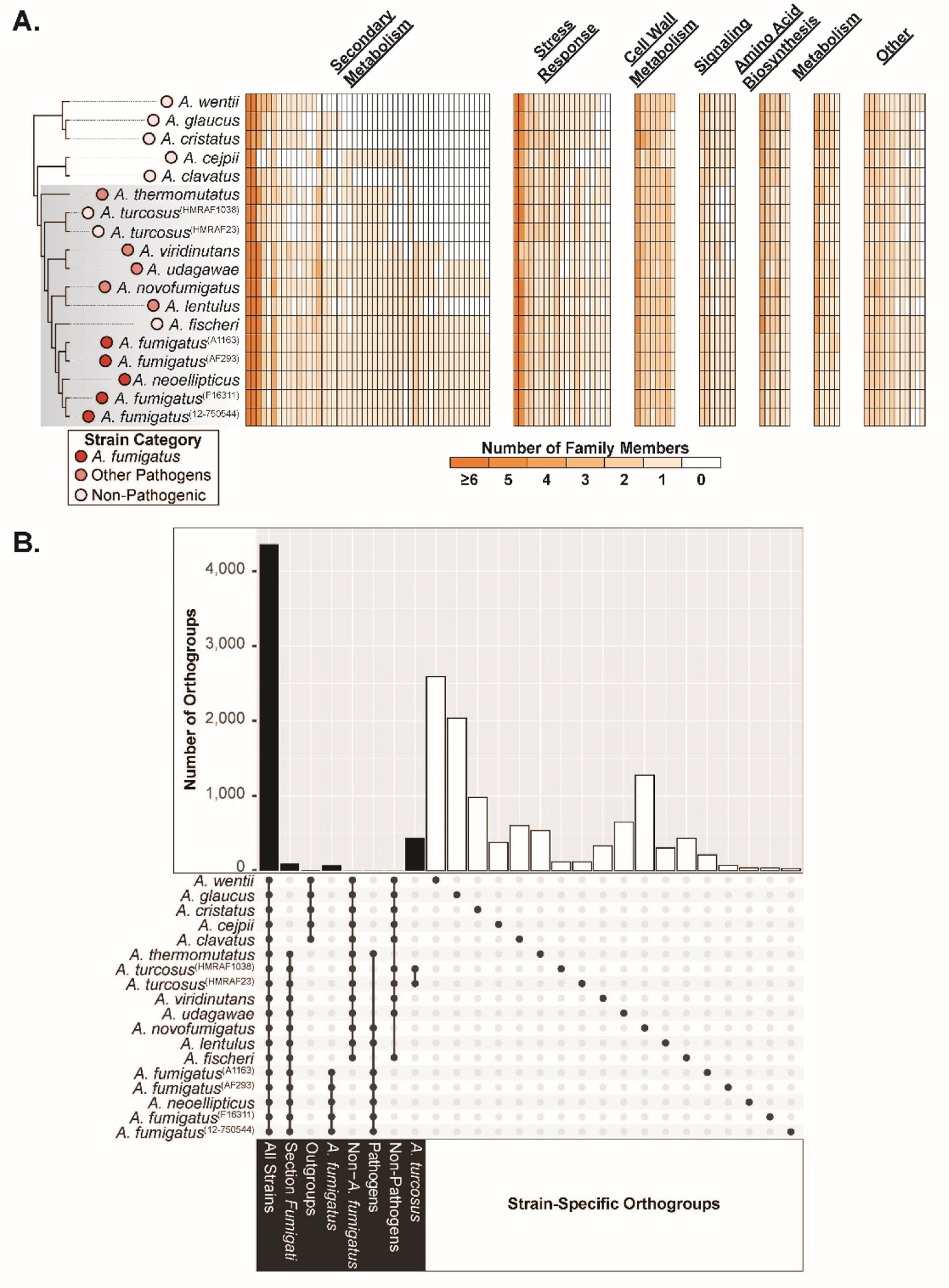
Gene families are largely conserved across section *Fumigati*, regardless of pathogenicity level. (A) Some virulence-related genes have different presence/absence patterns across strains in section *Fumigati.* Left, cladogram from Figure 1 showing the relationships amongst the 18 genomes studied. Gray box indicates strains belonging to section *Fumigati*. Right, heatmaps of the 105 / 189 gene families related to virulence that exhibited at least one gene presence/absence change in at least one species or strain, split into groups based on their general biological functions. Gene family labels can be found in Supplementary Table S3 in the same order presented here from left to right. (B) Gene families with representatives from all strains are the most prevalent. Upset plot (Conway et al. 2017) showing the number of all gene families present or absent in specific sets of strains. Black bars, gene family sets with members in more than one strain. White bars, strain-specific gene family sets.

We saw that 164 / 189 virulence-related gene families were already present in the last common ancestor of all section *Fumigati* species. Similarly, we estimated that on average, 12 genes have been lost from virulence-related gene families during the evolution of *A. fumigatus* strains 12-750544, F16311, and *A. neoellipticus* and 2 genes have been gained from virulence-related gene families during the evolution of *A. fumigatus* strains Af293 and A1163 compared to the *A. fumigatus* last common ancestor. The finding that many virulence-related genes are conserved across both pathogens and non-pathogens in section *Fumigati* suggests that most known genetic determinants of virulence likely evolved for functions other than causing disease in humans and have been instead recruited into pathogenic pathways in certain species.

Our unbiased approach consisted of analyzing the 14,294 gene families that resulted from constructing orthologous groups of genes from all 18 *Aspergillus* genomes. Similar to what we observed with virulence-related genes, we found that 4,361/14,294 gene families (∼31%) had family members in each of the 18 strains analyzed, and no gene families were present only in pathogens (Figure 2B). However, we found 98 gene families that were specific to section *Fumigati* (Figure 2B and Table S4). While the 98 gene families were not enriched for any Gene Ontology biological processes, molecular functions, or cellular compartments, the group contained genes associated with previously identified virulence-related traits, such as: a gene encoding a dimethylallyl tryptophan synthase (*cdpNPT* – <u>Afu8g00620</u>) located near the fumitremorgin-fumagillin-pseurotin supercluster (Yin et al. 2007; Wiemann et al. 2013), a major facilitator type transporter (*mdr3* – <u>Afu3g03500</u>) whose gene is highly expressed in *A. fumigatus* strains resistant to drugs (Nascimento et al. 2003; da Silva Ferreira et al. 2004), and a homolog of *mgtC* (<u>Afu7g05060</u>), a bacterial virulence factor required for survival in macrophages (Blanc-Potard 1997; Gastebois et al. 2011).

We found 72 gene families that were uniquely present in *A. fumigatus* (Figure 2B and Table S4). These *A. fumigatus*-specific genes were not enriched for any GO terms. We also found two gene families (predicted glucose-methanol-choline oxidoreductase family members - <u>NFIA_036190</u>/<u>NFIA_036210</u>, and a membrane dipeptidase - <u>NFIA_057190</u>) that had members in all strains except *A. fumigatus* and one gene family found only in non-pathogenic strains (a hypothetical protein with no identifiable domains – <u>NFIA_057720</u>). In summary, gene families are largely conserved across pathogens and non-pathogens in section *Fumigati,* but some genes whose possible role in virulence has not been tested were found only in *A. fumigatus*, thus supporting the species-specific pathogenicity model.

### Few gene families are associated with pathogenicity

Our gene family analysis did not identify genes whose presence/absence patterns were conserved in all pathogens found in section *Fumigati*. An alternative hypothesis is that the number of gene family members in a given strain could reflect the organisms’ ability to cause disease. To test this hypothesis we carried out a phylogenetically informed ANOVA (Revell 2012) on all 10,677 gene families that displayed a different number of gene family members in at least one taxon.

For this analysis we split the 18 *Aspergillus* taxa into three groups: *A. fumigatus*, other pathogens (that are not *A. fumigatus*), and non-pathogens. We found 83 gene families that had a statistically significant different number of members between groups (Figure S2). After conducting Tukey’s post-hoc test on all 83 gene families, we observed that almost all (72/83) of the gene families had more copies in *A. fumigatus* and were in fact those previously identified as “*A. fumigatus*-specific” in our strict gene presence/absence analysis (Table S4). One of the remaining 11 gene families was the membrane dipeptidase (<u>NFIA_057190</u>) found during our gene presence/absence analysis in all genomes other than *A. fumigatus*. The hypothetical gene family (<u>NFIA_057720</u>) found only in non-pathogenic species with the same gene presence/absence analysis (Figure 2B) was also identified via the phylogenetically informed ANOVA. The remaining nine gene families had the same number of genes in *A. fumigatus* and non-pathogens but a different number of family members in the other pathogens. Together, these data show that only very few gene families exhibit significant variation in their numbers across section *Fumigati* with respect to pathogenicity.

### Many genes experienced faster rates of evolution in pathogenic species

Another way in which genomes evolve that can affect phenotypes like virulence is through changes in the evolutionary rates of their constituent genes (Yang and Bielawski 2000). We carried out two evolutionary rate analyses to test whether genes in pathogens exhibit different rates of evolution compared to non-pathogens. For both analyses, our null hypothesis was that for a given single-copy gene family, a single rate (ω) represented the rate of sequence evolution for each gene family member in every strain, regardless of the pathogenicity level of the organisms examined (Figure 3A). In the first analysis, our alternative hypothesis was that each gene evolved at a unique rate in each of our three groups (*A. fumigatus* strains, other pathogens, and non-pathogens) (Figure 3Bi). We observed that 49% of all single-copy gene families (1,742/3,601) rejected the null hypothesis, suggesting that the evolutionary rate of these gene families differs among the three groups. Of the 1,742 gene families with three different ω values, 88% (1,532/1,742) had faster rates in pathogenic organisms (Figure 3Biii) and 10 had relatively high ω values (> 0.8) in *A. fumigatus* (Table S4). Each group also exhibited its own statistically different distribution of ω values (Figure 3Bii). Of the 81 / 189 virulence-related gene families that were present in a single copy in each *Aspergillus* genome, 56% (45/81) had exhibited different rates of evolution.

**Figure 3.**
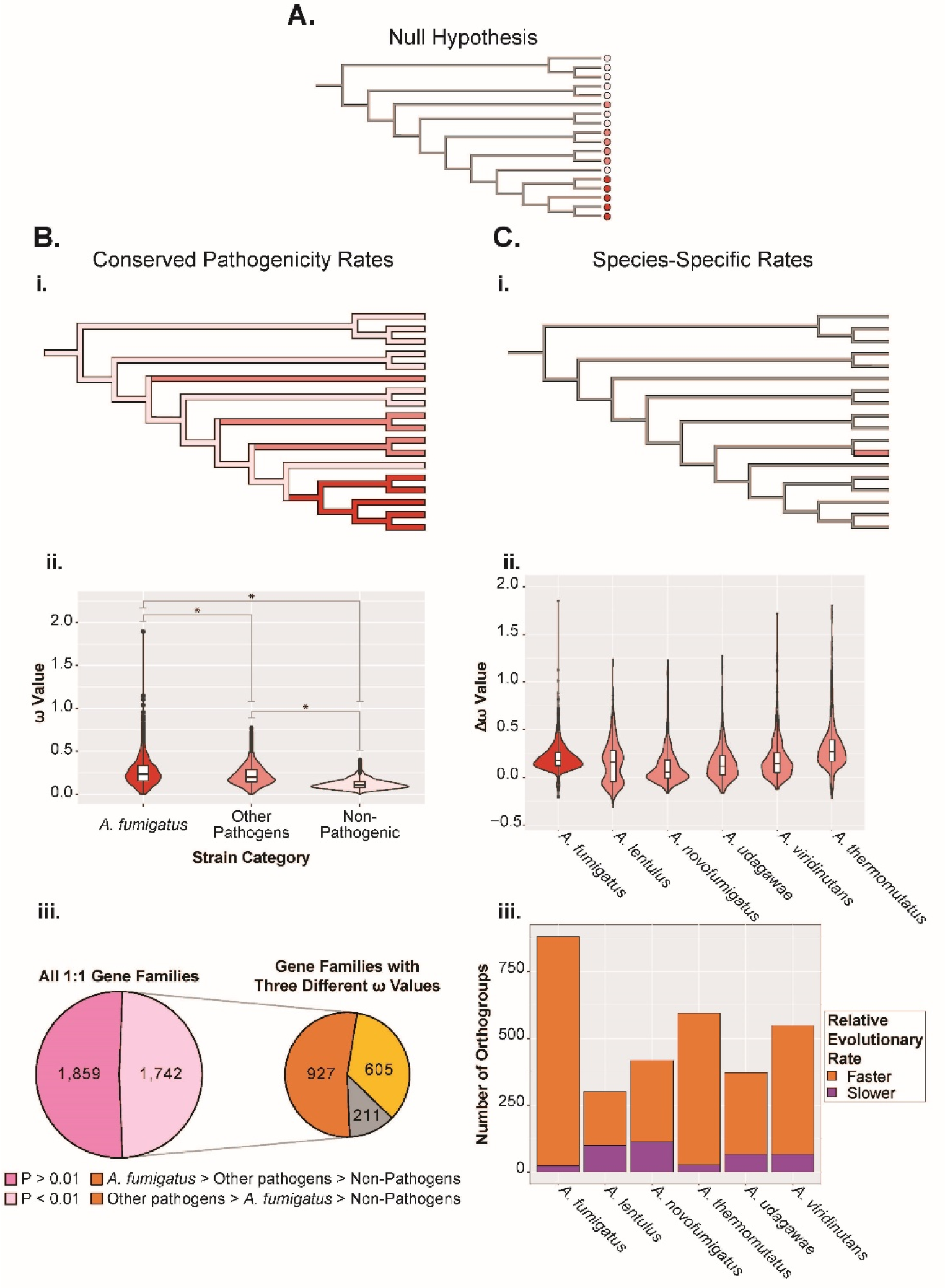
Genes in section *Fumigati* exhibit both pathogen- and species-specific rates of evolution. A) The null hypothesis that all branches in the phylogeny have the same ω value. (Bi) Alternative hypothesis examining the conserved pathogenicity model of gene evolution where genes are evolving at rates that correspond to their pathogenicity level. Branches are colored shades of red that correspond to their group (*A. fumigatus*, other pathogens, and non-pathogens). (Bii) Violin and box plots showing the ω values for each family member in 1,742 gene families (49% of all single-copy gene families) that exhibited different ω values (p-value <0.01) in the three groups of strains (*A. fumigatus*, other pathogens, and non-pathogens). Ten genes had ω values > 0.8 in highly pathogenic strains and are not shown here. *, adjusted p-value <0.0001 in a Paired Wilcoxon Signed-Rank test. (Biii) Left, pie chart showing the number of gene families that either did (light pink) or did not (dark pink) have different ω values for family members in strains with different pathogenicity levels. Right, pie chart further detailing the relative magnitude of ω values in strains with different pathogenicity levels. (Ci) A second alternative hypothesis examining the species-specific pathogenicity model of gene evolution where genes are evolving at one rate in one taxon and a different rate in all other taxa. Shown here is the model that was tested for *A. lentulus* where genes in *A. lentulus* experienced one rate of evolution, whereas their counterparts in all other taxa exhibited a different rate. (Cii) Violin and box plots showing the difference between ω values of genes in gene families whose family members had one ω in the pathogen and a different ω value for family members in other strains analyzed (p <0.01). ω value differences between the pathogen of interest and all other strains that were greater than two are not shown and constituted only 36/3,115 comparisons that exhibited p<0.01. Strains are colored the same as in panel B. (Ciii) Bar chart showing the number of gene families where the family member in the pathogen is evolving faster or slower than the copies in the other strains. Taxa in all phylogenies are in the same order as in Figure 1.

In the second analysis, we tested if each gene was evolving at a unique rate in each pathogen, relative to the other species we analyzed (Figure 3Ci). We found that, on average, 519 / 3,601 single-copy genes exhibited a different rate in the pathogen of interest than in the rest of species, with most evolving faster in the pathogens (Figure 3Cii and 3Ciii). *A. fumigatus* had the highest number of genes whose rates differed from the rest of the species (881), while *A. lentulus* had the fewest (301) (Figure 3Ciii and Table S4). Of the 881 genes whose evolutionary rate differed between *A. fumigatus* and the rest of the taxa, only 34 were not also included among the 1,742 genes whose evolutionary rates differed between pathogens and non-pathogens (Table S4). Overall, our data show that genes in pathogens are evolving faster than in non-pathogens, both in a conserved and species-specific manner.

### Transcription factors with pathogenicity-related patterns of evolution have diverse effects on virulence

To test if any of the genes whose evolutionary signatures differed between pathogens and non-pathogens directly affected either fungal or host survival, we tested 17 knockout strains of transcription factor-encoding (TF) genes in two virulence-related assays. One TF (<u>Afu7g00210</u>) was found only in *A. fumigatus* (Figure 2B), one (<u>Afu6g08540</u>) was identified as being fast-evolving in *A. fumigatus*, four (<u>Afu2g17895</u>, <u>Afu3g02160</u>, <u>Afu7g04890</u>, and *gliZ* - <u>Afu6g09630</u>) were members of gene families with a non-statistically significant higher number of family members in pathogens, five (<u>Afu1g11000</u>, <u>Afu2g00470</u>, <u>Afu6g11750</u>, <u>Afu3g00210</u>, and <u>Afu8g05750</u>) were members of gene families with a non-statistically significant higher number of family members in *A. fumigatus*, and six (<u>Afu2g17860</u>, <u>Afu5g01065</u>, <u>Afu5g14530</u>, <u>Afu1g01340</u>, <u>Afu3g01640</u>, and <u>Afu2g16310</u>) were found only section *Fumigati*. None of the TF mutants exhibited a growth defect compared to their parent strain (CEA17) when grown in conventional lab conditions (Figure S3).

In the first assay, asexual spores (conidia) or germlings from either a background strain of *A. fumigatus* (CEA17) or one of the *A. fumigatus* knockout mutants of transcription factors (Furukawa et al. 2020) were incubated with *Protostelium aurantium,* a fungivorous amoeba used to study how fungi may have evolved the ability to evade or survive phagocytosis by human immune cells (Radosa et al. 2019). We found that mutant asexual spores of four genes (<u>Afu2g17860</u>, <u>Afu2g17895</u>, <u>Afu7g04890</u>, and <u>Afu2g00470</u>) exhibited an increase in survival relative to CEA17 (Figure 4A). In contrast, while germlings of many mutants showed a qualitative difference in survival relative to CEA17, those differences were not statistically significant (Dunn’s Test adjusted p-value > 0.1).

**Figure 4.**
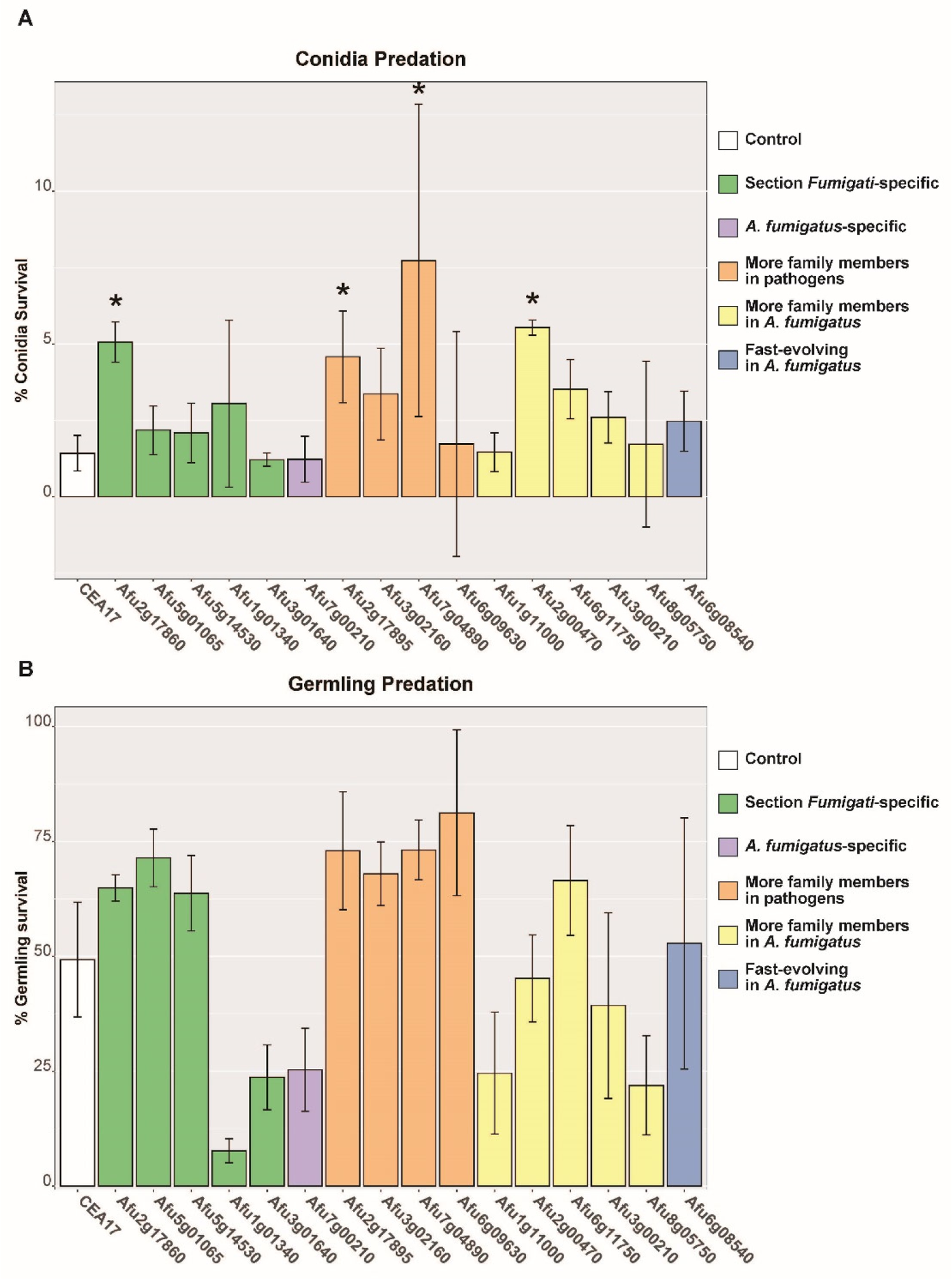
Multiple transcription factors with virulence-related genomic traits affect the survival of *A. fumigatus* during amoeba predation. (A) Survival of swollen *A. fumigatus* asexual spores (conidia) after interaction with *Protostelium aurantium*. Spores of *A. fumigatus* were incubated 4 hours at 37°C in Czapek-Dox medium and confronted with *P. aurantium* at a prey-predator ratio of 10:1 (10^5^ spores and 10^4^ trophozoites of *P. aurantium*). Survival is expressed as the relative reduction in the metabolic rate of the fungus in comparison to amoeba-free controls over three hours. Data represent the mean and standard deviation of three biological replicates. *, p<0.1 in an adjusted Dunn Test comparing the survival of the mutant strain to the parental strain CEA17. (B) Survival of *A. fumigatus* germlings after interaction with *P. aurantium*. Asexual spores of *A. fumigatus* were pre-grown to germlings for 10 hours at 37°C in Czapek-Dox medium and confronted with *P. aurantium*. All other assay parameters are the same as in panel A. No mutant strain exhibited statistically significant difference in survival relative to CEA17 in an adjusted Dunn Test. Both asexual spores and germling confrontation assays were confirmed to have significant p-values (<0.05) in Kruskal-Wallis tests before carrying out the post-hoc test and mutant strains did not exhibit large growth phenotypes in the absence of amoeba (Figure S4). Mutants are color-coded based on genomic traits related to pathogenicity that they possess. Knockout mutants were constructed in the CEA17 background (Furukawa et al. 2020), but *A. fumigatus* strain Af293 gene ids for the corresponding orthologous genes are shown here and in Figure 5. Gene absence patterns were confirmed with tblastn. Note that a potential, low-confidence ortholog of Afu5g01065 was found in *A. wentii*; however, all other *A. fumigatus* genes were found missing in the species listed. The mutant of Afu2g16310 could not be assayed due to technical reasons.

In the second assay, we measured virulence in the greater wax moth (*Galleria mellonella*) model of *A. fumigatus* disease. We found that almost one third of all knockout mutants tested (5 / 17) exhibited a statistically significant decrease in virulence (Figures 5 and S5). One of the transcription factor mutants that resulted in a significant decrease in larval killing was that of *gliZ* (<u>Afu6g09630</u>), a regulator of gliotoxin production (Bok et al. 2006) that we observed was found in all pathogenic strains but missing in all non-pathogenic species except *A. cejpii* and *A. fischeri*. To our knowledge, this is the first time *gliZ* has been tested in the greater wax model of fungal disease or shown to contribute to fungal pathogenesis. In addition to *gliZ*, another of the knockout mutants with a statistically significant decrease in virulence also supported the conserved pathogenicity model as the gene knocked out in that strain (<u>Afu7g04890</u>) was also in a gene family that in general has more family members in pathogens. The other three knockout mutants that were less virulent supported the species-specific pathogenicity model as the genes knocked out in two of them (<u>Afu3g00210</u> and <u>Afu8g05750</u>) were in gene families that in general had more family members in *A. fumigatus* and gene knocked out in the other mutant (<u>Afu7g00210</u>) was only found in *A. fumigatus*. These results suggest that genes whose evolution differs between pathogens and non-pathogens are likely to contribute to disease-related traits.

**Figure 5.**
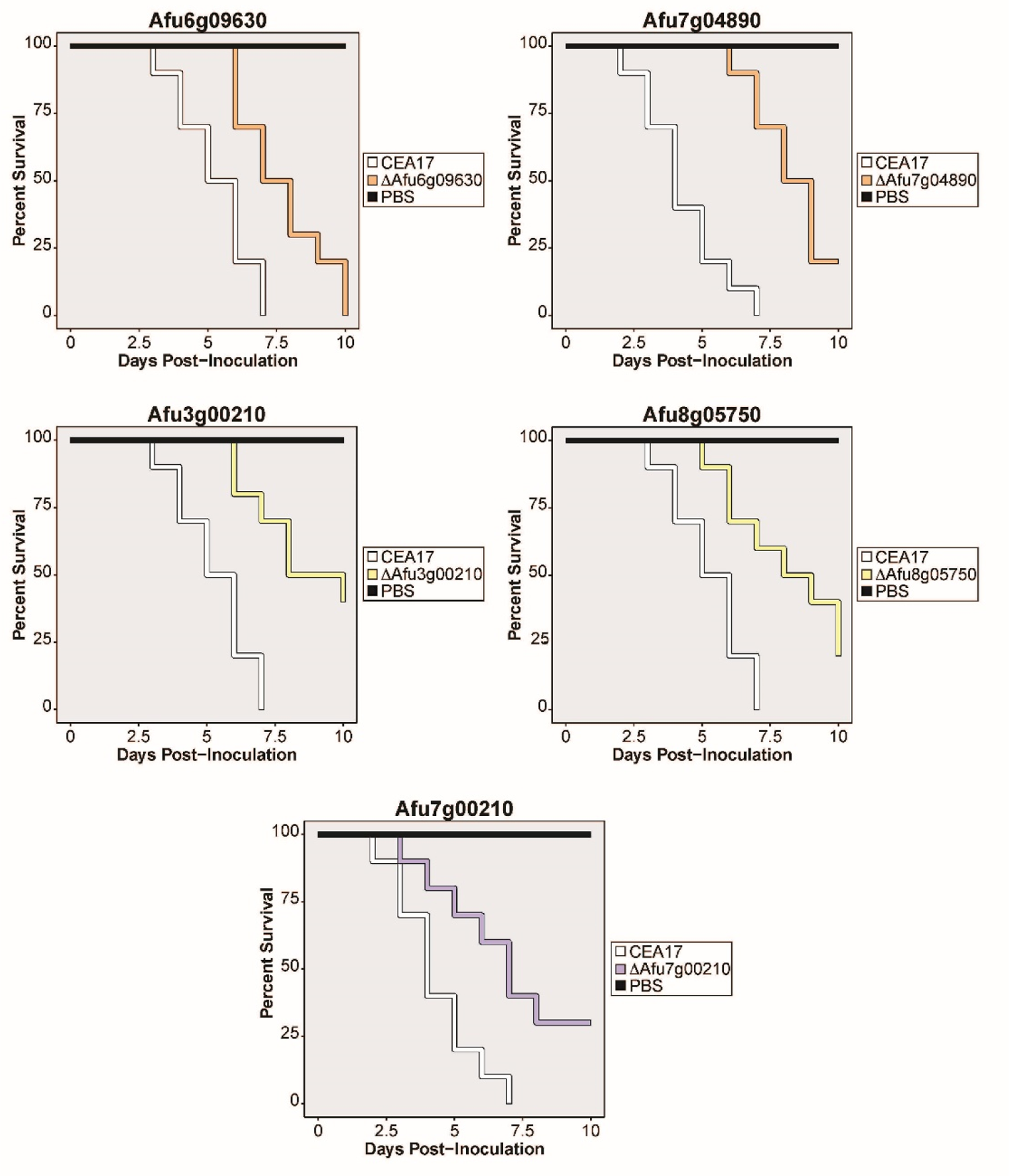
Multiple transcription factors in *A. fumigatus* whose evolution differs with respect to pathogenicity affect virulence in the greater wax moth model of disease. Cumulative survival of *Galleria mellonella* larvae inoculated with phosphate buffered saline (Black), asexual spores of the parental strain CEA17 (white), and asexual spores from null mutants of transcription factors whose evolution differs with respect to pathogenicity (various colors). Ten larvae were used per inoculation in all assays. Color scheme is the same as in Figure 4. All mutant survival curves shown here were statistically different (p<0.008 in a Log-Rank test) from the CEA17 survival curve. Mutants whose survival curves are shown in orange have significantly more gene family members in pathogens, those in yellow have significantly more family members in *A. fumigatus*, and the mutant in purple is *A. fumigatus*-specific. Note that the results of the mutants in orange support the conserved pathogenicity model, whereas those of the mutants in yellow and purple support the species-specific pathogenicity model. The Afu6g09630 gene is *gliZ*, a regulator of the biosynthesis of the secondary metabolite gliotoxin, a known modulator of host biology.

## Discussion

Our results show that some genomic traits related to virulence are shared amongst all pathogens in section *Fumigati* (supporting the “conserved pathogenicity” model), others are found only in individual pathogenic species or strains (supporting the “species-specific pathogenicity” model), and previously described virulence-related genes likely function in a context-dependent manner as they are largely present in both pathogens and non-pathogens (Figure 2A). Giving support for the species-specific model, our gene presence / absence analyses across the 18 strains showed that there were 72 *A. fumigatus*-specific gene families, but no pathogen-specific gene families (Figure 2B). However, our evolutionary rate analyses offered more support for the conserved pathogenicity model as over 1,700 gene families showed pathogen-specific evolutionary rates and only 34 gene families exhibited evolutionary rates specific to *A. fumigatus* (Figure 3).

Multiple transcription factors we identified as having virulence-related genomic traits also displayed roles in diverse virulence-related assays both here (Figure 4 and 5) and in previously published studies of *A. fumigatus* virulence. Of the eight transcription factors whose null mutants exhibited at least one phenotype in our two assays, half of them were downregulated and none were upregulated during the switch to human body temperature (Lind et al. 2016). Additionally, *gliZ*, a regulator of gliotoxin production whose gene family we found to be largely pathogen-specific and whose mutant was less virulent than the WT strain, was heavily upregulated in *A*. *fumigatus* germlings that were extracted 12-14 hours after mouse infection (McDonagh et al. 2008) while the seven other TFs we studied were not differentially regulated during the early events of mouse infection. Taken together, our functional assays show that our evolutionary genomic approach has power to identify genes both previously connected to *A. fumigatus* virulence and novel ones.

We analyzed all sequenced species in section *Fumigati* (as of July 2019) and a representative sampling of strains from *A. fumigatus*, carried out a diverse set of evolutionary genomic analyses, and functionally tested our identified genes in multiple assays, thus building on previous studies that used smaller numbers of section *Fumigati* species and close relatives in *Aspergillus* and focused on strict gene presence / absence (Fedorova et al. 2008). Previous work also compared *A. novofumigatus*, one of the section *Fumigati* species we considered here, to its relative *A. fumigatus* (Kjærbølling et al. 2018), and while that study used a broader and less stringent list of virulence-related genes that also included allergens, they also saw high levels of gene conservation between the two species. Together, the two studies support the hypothesis that *A. novofumigatus* could be nearly as pathogenic as *A. fumigatus* due to this conservation of almost all virulence-related genes. In section *Flavi*, another taxonomic section in genus *Aspergillus* that contains the human and plant pathogen *Aspergillus flavus*, it has been hypothesized that transcription factors may be linked to pathogenicity (Kjærbølling et al. 2020), and similarly, we saw that one of the 72 *A. fumigatus*-specific genes we identified is a transcription factor that contributes to disease in *Galleria mellonella* larvae (Figure 5).

As more genomes from strains and species in section *Fumigati* become available, our power to detect quantitative differences will increase, and allow us to more robustly test the conserved pathogenicity model and expand our species-specific pathogenicity model to include “strain-specific” elements. This will be especially important considering the continued and growing appreciation for strain-specific traits and differences in *Aspergillus* genomes and pathogenicity (Keller 2017; Bastos et al. 2020; Dos Santos et al. 2020; Steenwyk, Lind, Ries, et al. 2020; Kowalski et al. 2021). Future studies will also place *A. oerlinghausenensis*, another species closely related to *A. fumigatus* (Houbraken et al. 2016), within this evolutionary framework of pathogenicity, but due to our previous genome-wide phylogenomic analyses of *A. oerlinghausenensis*, *A. fischeri*, and *A. fumigatus* (Steenwyk, Mead, Knowles, et al. 2020), we do not anticipate that inclusion of *A. oerlinghausenensis* will drastically change our findings.

Evolutionary studies have also been caried out in fungal pathogens outside of the genus *Aspergillus*, and when our results are placed in the context of this literature, a diverse set of mechanisms have driven the evolution of fungal pathogenicity. The ability to infect humans has also evolved multiple times in *Candida* species found within the fungal subphylum Saccharomycotina (Gabaldón et al. 2016); however, gene family expansion and interspecies hybridization were much more important for the evolution of pathogenicity in that clade compared to the results we present here in section *Fumigati* where there was little evidence of dramatic changes in gene family member number between pathogens and non-pathogens (Figure S2). Similarly, gene family size was hypothesized to be an important factor in the evolution of *Coccidioides* pathogens, and just as we only saw 84 genes with *A. fumigatus*-specific evolutionary rates, this group of pathogens had a relatively small number genes with species-specific evolutionary rates (Sharpton et al. 2009). In pathogenic *Cryptococcus* species, mating type loci and the switch from a tetrapolar to bipolar mating system have been suggested as being key in producing the genomic environment necessary for pathogenicity to evolve (Sun et al. 2019), but in *A. fumigatus*, mating type loci do not appear to contribute to virulence (Losada et al. 2015) and the contribution of mating across section *Fumigati* has only rarely been studied (Rydholm et al. 2007).

Worldwide mortality rates for aspergillosis infections are estimated to range from as high as 95% to as low as 30%, and drug resistance is a frequent worry for clinicians (Brown et al. 2012). To combat this global health issue, more must be understood about *Aspergillus* biology and evolution. Here, we have presented many promising, novel candidates for future study and have placed them within an evolutionary context that will also guide their study with relation to non-*A. fumigatus* pathogenic species found within section *Fumigati*. Our data infer the evolution of pathogenic and non-pathogenic *Aspergillus* genomes and provide clues on how *Aspergillus* pathogenicity evolved. Furthermore, genes that fit the conserved pathogenicity model may be useful as targets for the treatment of disease caused by all section *Fumigati* species, whereas genes that fit the species-specific pathogenicity model may be useful for species-specific treatment strategies. More generally, this work provides the basis for an evolutionary framework that can inform multiple aspects of the study of both *Aspergillus* species and the diseases they cause.

## Methods

### Genome procurement, assembly, and annotation

Genomes and annotations for *A. fumigatus* strains Af293 and A1163, along with all non-*A. fumigatus*, publicly available (as of July 2019) annotated genomes from section *Fumigati* were downloaded for analyses. We also obtained genomes and annotations for four outgroup species to facilitate phylogenetic analyses and comparisons. To expand the number of genomes analyzed, we assembled and/or annotated five additional *Aspergillus* genomes. More specifically, raw genomic reads for *A. fumigatus* strains F16311 and 12-7505446 were downloaded from NCBI for genome assembly and annotation. These strains were chosen because they, together with *A. fumigatus* strains Af293 and A1163, span the known diversity of *A. fumigatus* (Lind et al. 2017); additionally, available genomes for *A. cejpii* FS110, *A. neoellipticus* NRRL 5109, and *A. viridinutans* FRR 0576 were downloaded from NCBI and annotated (Abdolrasouli et al. 2015; Li et al. 2018; Urquhart et al. 2019) (Table S1). To quality trim sequence reads, we used Trimmomatic, version 0.36 (Bolger et al. 2014) using parameters described elsewhere (Steenwyk and Rokas 2017). The resulting high-quality reads were used as input to the genome assembly software SPAdes, version 3.8.1 (Bankevich et al. 2012), with the ‘careful’ parameter to reduce mismatches and short indels and the ‘cov-cutoff’ parameter set to ‘auto.’ Partial and complete gene models were predicted using Augustus, version 2.5.5 (Stanke and Waack 2003), with the ‘minexonintronprob’ and ‘minmeanexonintronprob’ parameters set to 0.1 and 0.4, respectively. Genome annotation quality was assessed using BUSCO, version 2.0.1 (Waterhouse et al. 2018), with the Pezizomycotina database of orthologs from OrthoDB, version 9 (Waterhouse et al. 2013). Genome annotation quality was similar between publicly available genomes and genomes assembled and/or annotated in the present project. For example, the publicly available assembly and annotation for *A. fumigatus* strain A1163 had 94.0% of BUSCO genes present in single copy while the assembled and annotated genome for *A. fumigatus* strain F16311 had 93.9% of BUSCO genes present in a single copy.

### Phylogenomic data matrix construction and analyses

We employed a maximum likelihood framework to reconstruct the evolutionary history of the 18 *Aspergillus* taxa. We first identified single-copy orthologous genes by clustering genes with high sequence similarity into orthologous groups using Markov clustering (van Dongen 2000) as implemented in OrthoMCL, version 1.4 (Li et al. 2003), with an inflation parameter of 2.8. Gene sequence similarity was determined using a blastp “all-vs-all” using NCBI’s Blast+, version 2.3.0 (Camacho et al. 2009) with an e-value cutoff of 1e-10, a 30% identity cutoff, and a 70% match cutoff. Out of a resulting 14,294 orthologous groups, 3,601 orthologous groups had all 18 taxa represented by a single sequence which, are hereafter referred to as single-copy orthologous genes. The protein sequences of the 3,601 single-copy orthologous genes were individually aligned using Mafft, version 7.402 (Katoh and Standley 2013), using the same parameters as described elsewhere (Steenwyk, Shen, et al. 2019). Nucleotide sequences were threaded onto the protein alignments using the thread_dna function in PhyKIT, version 0.1 (Steenwyk et al. 2021). The codon-based sequences were subsequently trimmed using trimAl, version 1.2rev59 (Capella-Gutierrez et al. 2009), using the ‘automated1’ parameter. The resulting single-gene alignments were concatenated into a single data matrix using the create_concat function in PhyKIT, version 0.1 (Steenwyk et al. 2021).

To infer the evolutionary history of *Aspergillus* species in section *Fumigati* and the outgroup taxa, we implemented a concatenation without gene-based partitioning, concatenation with gene-based partitioning, and gene-based coalescence (Felsenstein 1981; Rokas et al. 2003; Edwards 2009; Zhang et al. 2018). For concatenation without gene-based partitioning, we used the 3,601-gene matrix as input to IQ-TREE (Nguyen et al. 2015) and inferred the best-fitting model of substitutions according to Bayesian information criterion values using the “-m TEST” parameter. The best-fitting model was determined to be a general time-reversal model with invariable sites, empirical nucleotide frequencies, and a discrete gamma model with four rate categories or “GTR+F+I+G4” (Tavaré 1986; Yang 1994; Gu et al. 1995). Lastly, we increased the number of candidate trees used during maximum likelihood search by setting the “-nbest” parameter to 10. Bipartition support was assessed using 5,000 ultrafast bootstrap approximations (Hoang et al. 2018). We refer to the tree inferred using this method as the reference tree topology depicted in Figure 1.

To infer the evolutionary history of *Aspergillus* species in section *Fumigati* and the outgroup strains using concatenation with gene-based partitioning and coalescence, we first determined the best-fitting model of substitution using the “-m TEST” parameter and reconstructed the phylogeny of the 3,601 single-copy orthologous genes individually using default IQ-TREE parameters (Nguyen et al. 2015). For concatenation with gene-based partitioning, we created a nexus-format partition file that describes gene boundaries in the 3,601-gene matrix and the best-fitting model of substitutions for each partition. We used the nexus-format partition file as input using the “-spp” parameter along with the concatenated 3,601-gene matrix to reconstruct the Fumigati phylogeny. Bipartition support was assessed using 5,000 ultrafast bootstrap approximations (Hoang et al. 2018). For coalescence, we first collapsed lowly supported bipartitions in all single-gene trees defined as less than 80% ultrafast bootstrap approximation support to reduce signal from poorly supported bipartitions. To do so, we assessed bipartition support using 5,000 ultrafast bootstrap approximations for individual single-gene trees (Hoang et al. 2018). To infer a coalescence-based phylogeny, we combined all single-gene trees with collapsed bipartitions into a single file and used it as input to ASTRAL-III, version 5.6.3 (Zhang et al. 2018), with default parameters. Bipartition support was assessed using posterior probabilities.

### Gene Ontology Enrichment Analyses

To determine if lists of genes of interest contained enriched Gene Ontology terms, we used GOATOOLs version 0.9.7 (Klopfenstein et al. 2018). Annotations for the *A. fumigatus* Af293 genome were downloaded from version 45 of FungiDB (Basenko et al. 2018), and the 2019-07-01 version of the basic Gene Ontology (Ashburner et al. 2000; The Gene Ontology Consortium et al. 2021) was used for all analyses. A term was considered enriched if it had an adjusted p-value (using the Benjamini-Hochberg method) less than 0.05.

### Gene Family Expansions and Contractions

To study if the number of gene family members is expanded or contracted in classes of strains (pathogens or non-pathogens) or specific strains, we carried out a phylogenetically-informed analysis of variance with the phylANOVA function located within version 0.7-70 of the phytools package (Revell 2012). Taxon relationships were provided from the phylogenetic tree that resulted from the concatenation without gene based partitioning approach, and only those gene families that had at least one change in gene family number were used in the analysis. The simulation-based ANOVA was performed for each gene family and run with 10,000 simulations in order to derive a p-value reflecting if the average number of genes was different in the three groups of strains (*A. fumigatus*, other pathogens, and non-pathogens). P-values were then corrected using the Benjamini-Hochberg method found within the “p.adjust” function in R (R Core Team 2018). Gene families were considered significantly different if their adjusted p-values were less than 0.05. Tukey’s range post-hoc test from the Python module “statsmodels” version 0.10.0 (Seabold and Perktold 2010) was then carried out on significantly different gene families in order to determine if the average number of gene family members differed in any of the pairwise comparison (ex. the number of genes in *A. fumigatus* vs non-pathogenic species).

### Gene family history and rates of molecular evolution

To determine the evolutionary history of gene families across section *Fumigati* species and outgroups, we implemented a maximum likelihood framework with a birth-death innovation model and gamma-distributed rates across families as implemented in DupliPHY-ML (Ames et al. 2012). DupliPHY-ML takes as input a matrix of gene family copy number and a phylogeny. To construct a matrix of gene family copy number, we used orthologous groups of genes as proxies for gene families and used the number gene sequences for a given species as the copy number information per gene family. For the phylogeny, we used the reference phylogeny described previously.

To determine the rate of sequence evolution across the evolutionary history of *Fumigati* species on a per gene basis, we used measures of the rate of nonsynonymous substitutions (dN) over the rate of synonymous substitutions (dS) (hereafter referred to as dN/dS or ω) using an approach described elsewhere (Steenwyk, Opulente, et al. 2019). To do so, we used untrimmed codon-based alignments generated during the construction of the 3,601-gene matrix used for phylogenomic analyses. For each of the 3,601 genes, we calculated ω using PAML, version 4.9 (Yang 2007), under two hypotheses: a null hypothesis (H_O_) and an alternative hypothesis (H_A_). For H_O_, we allowed a single ω value to represent the rate of sequence evolution across the reference phylogeny. For the first H_A_, we tested if different groups (*A. fumigatus*, other pathogens, or non-pathogens) were associated with different rates of sequence evolution. For the second H_A_, we tested if each gene was evolving at a unique rate in each pathogen, relative to the other branches in the tree (Figure 3Ci). For each comparison, to determine if H_A_ significantly differed from H_O_, we used a likelihood ratio test (α = 0.01).

### Amoeba Predation Assays

Asexual spores (conidia) of *A. fumigatus* (either transcription factor mutants previously described (Furukawa et al. 2020) and obtained as described in (Zhao et al. 2019) or the background strain CEA17) were incubated 4 h at 37°C in Czapek-Dox medium (CZD, Sigma-Aldrich Chemie, Munich, Germany) to induce swelling, and confronted with *Protostelium aurantium* at a prey-predator ratio of 10:1 (10^5^ conidia and 10^4^ trophozoites of *P. aurantium*) for 18 h at 22°C. Afterwards, the assay plate was incubated for 1 h at 37°C to inactivate the amoebae. Subsequently, 0.002% [w/v] resazurin (Sigma-Aldrich, Taufkirchen, Germany) was added and metabolic rates were calculated from the time dependent reduction of resazurin to the fluorescent resorufin over 3 h at 37°C using an Infinite M200 Pro fluorescence plate reader (Tecan, Männedorf, Switzerland). Survival was determined from the difference in the metabolic rates of the fungus after amoeba confrontation and amoeba-free controls. These controls were also used to determine the fitness of each strain in CZD-medium. Essentially the same assay was carried out to determine the survival of germlings of *A. fumigatus*, except that conidia of *A. fumigatus* were pre-grown to germlings for 10 h at 37°C in CZD medium before the addition of trophozoites of *P. aurantium*.

### Virulence assays in the great wax moth (*Galleria mellonella*) model of fungal disease

*Galleria mellonella* larvae were obtained by breeding adult larvae (Fuchs et al. 2010) weighing 275-330 mg in starvation conditions in petri dishes at 37°C in the dark for 24 hours prior to infection. All selected larvae were in the final stage of larval (sixth) stage development. Fresh conidia of each strain of *A. fumigatus* were obtained. For each strain, spores were counted using a hemocytometer and the initial concentration of the spore suspensions for the infections were 2×10^8^ conidia/ml. A total of 5 μl (1×10^6^ conidia/larvae) of each suspension was inoculated per larva. The control group was composed of larvae inoculated with 5 μl of PBS to observe death by physical trauma. The inoculation was performed using a Hamilton syringe (7000.5KH) through the last left proleg. After infection, the larvae were kept at 37°C petri dishes in the dark and scored daily. Larvae were considered dead when a lack of movement was observed in response to touch. The viability of the inoculum administered was determined by plating a serial dilution of the conidia in 37% YAG medium. The statistical significance of the comparative survival values was calculated using the log rank analysis of Mantel-Cox and Gehan-Brestow-Wilcoxon found in the statistical analysis package Prism.

## Acknowledgements

We thank members of the Rokas lab for enlightening discussions and comments on the project, especially Dr. Xing-Xing Shen and Dr. Abigail LaBella. We also thank Dr. Xing-Xing for conducting preliminary evolutionary analyses that were not included in this manuscript.

