## Supplementary Figures for "An evolutionary genomic approach reveals both conserved and species-specific genetic elements related to human disease in closely related *Aspergillus* fungi"

Short title: Evolution of *Aspergillus* pathogenicity

* MEM and JLS contributed equally to this work

Key Words: evolution, *Aspergillus fumigatus*, evolutionary rate, virulence, pathogenicity, fungal disease, aspergillosis, convergent evolution

#### Supplementary Figures

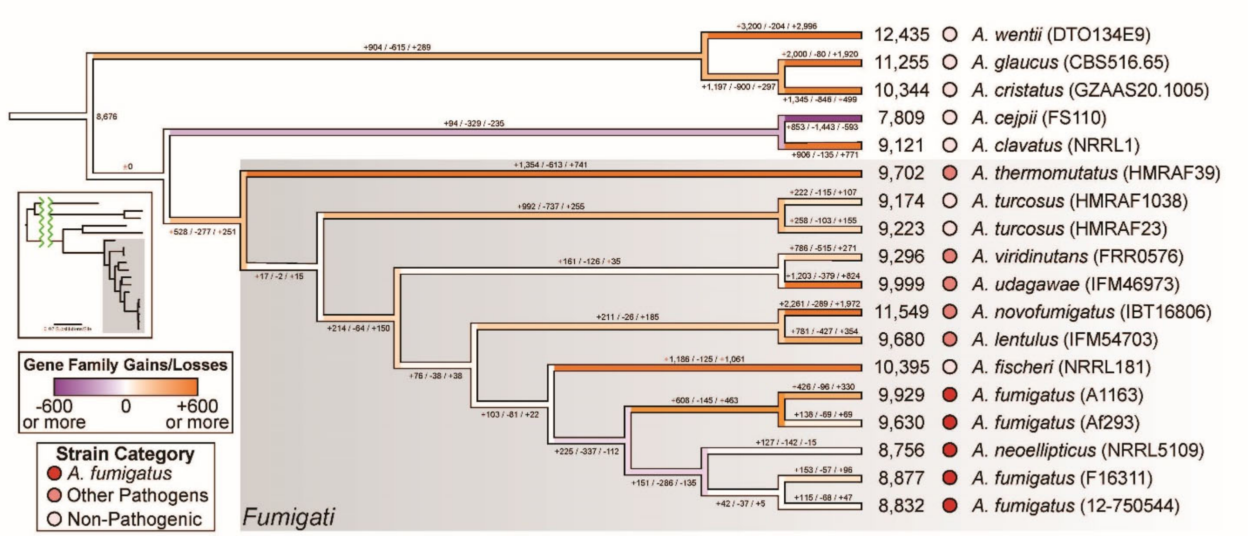


**Supplementary Figure S1. Genome-scale phylogeny of select genomes from section *Fumigati* showing gene family gains/losses.** Relationships among taxa in section *Fumigati* inferred from a concatenation-based, maximum likelihood approach. The number of gene family gains, losses, and the net gain or less (in that order) is shown above branches. Branches are colored based on the number of net gene family gains or losses. Numbers at branch tips represent the total number of genes in that genome. Pathogenicity level (high, middle, and low) of each strain is shown via shades of red in circles and is based on approximately how frequently that species is found in the clinic. Strain designations are in parenthesis next to species names. Insert tree shows the phylogeny with branch lengths reflective of the estimated number of nucleotide substitutions per site and taxa in the same order as the larger cladogram.

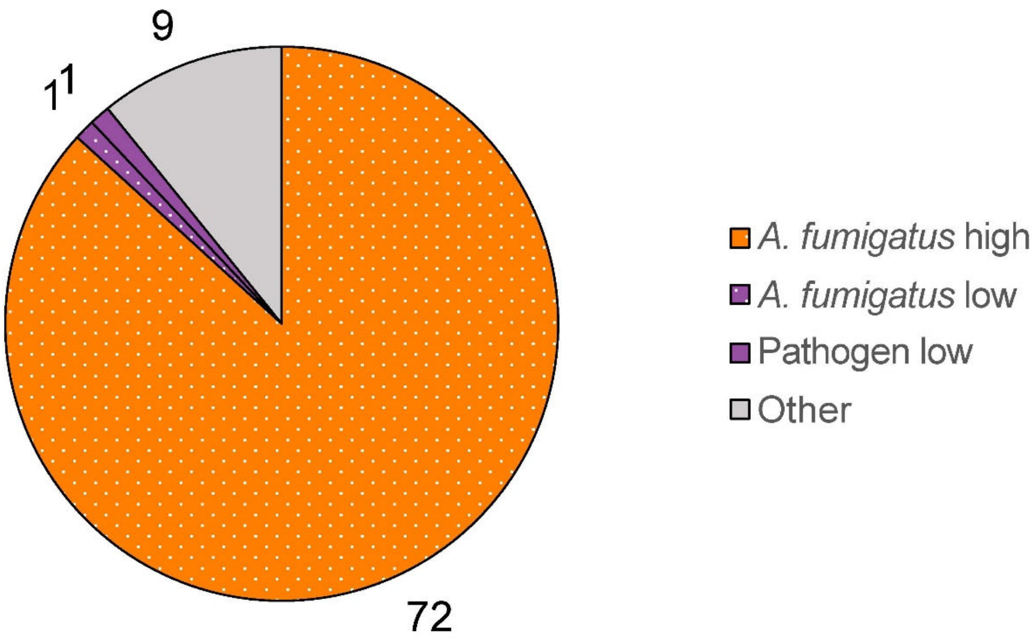


**Supplementary Figure S2. The majority of orthogroups with different numbers of family members in *A. fumigatus*, other pathogens, and non-pathogens are *A. fumigatus*-specific.** Pie chart showing the number of orthogroups with a statistically significant different number of family members in *A. fumigatus* or groups of strains relative to all other strains studied. "*A. fumigatus* high", more family members in *A. fumigatus* strains compared to all other strains. "*A. fumigatus* low", fewer family members in *A. fumigatus* strains compared to all other strains. "Pathogen low", fewer family members in A. fumigatus and other pathogenic strains compared to the non-pathogens. "Other", more family members in the non-*A. fumigatus*, pathogenic strains compared to all other strains.

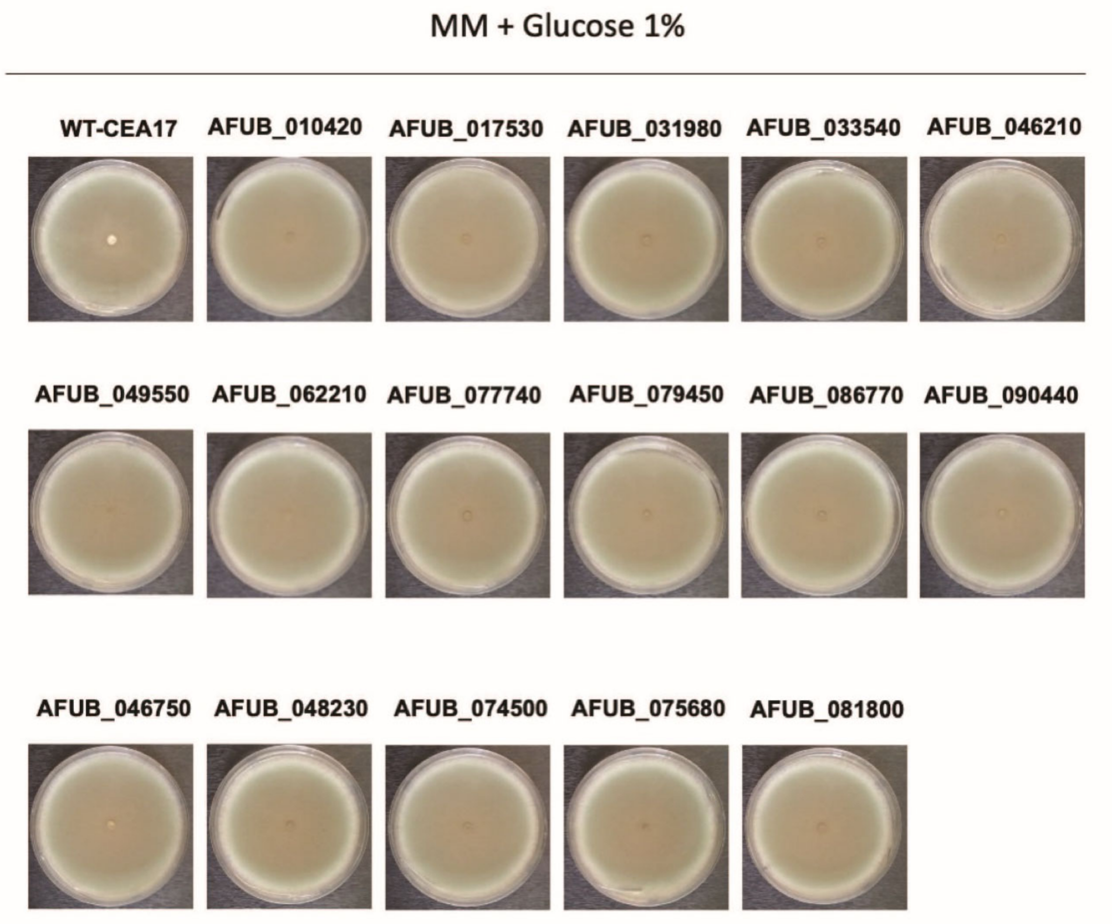


**Supplementary Figure S3: Transcription factor mutants do not have growth defects when grown under normal lab conditions.** Transcription factor mutants and the background strain CEA17 were grown in Aspergillus Minimal Media supplemented with 1% Glucose. Image IDs are for the A1163 orthologs of the AF293 IDs given in Figures 4 and 5.

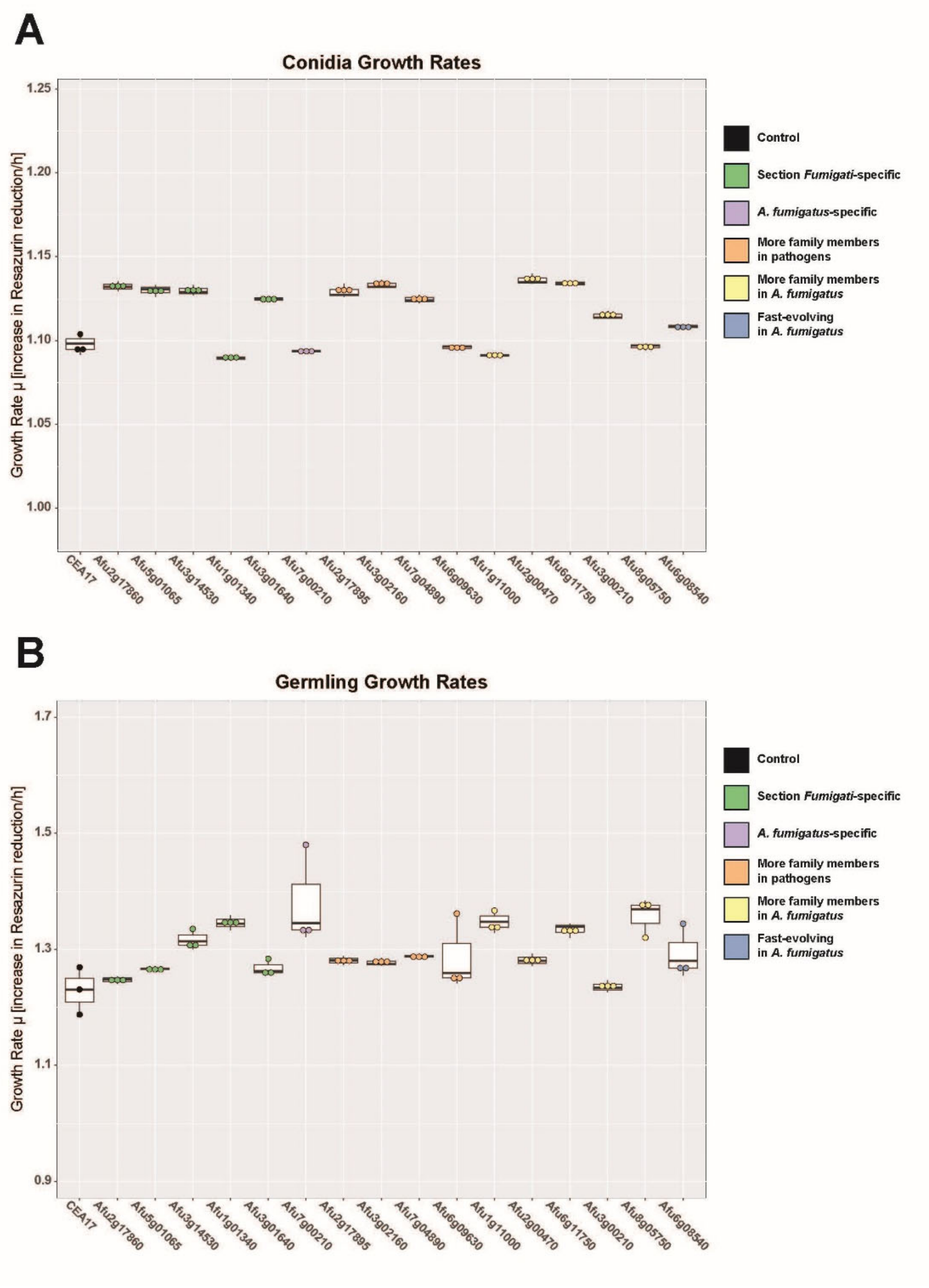


**Supplementary Figure S4. Amoeba predation growth assay controls.** Asexual spores or conidia (A) or germlings (B) of *A. fumigatus* were grown in Czapek-Dox medium and the metabolic rate was determined over a growth period of three hours. The metabolic rates were calculated form the time-dependent, exponential increase in resazurin reduction to the fluorescent resorufin. Box plots show the results of three biological replicates.

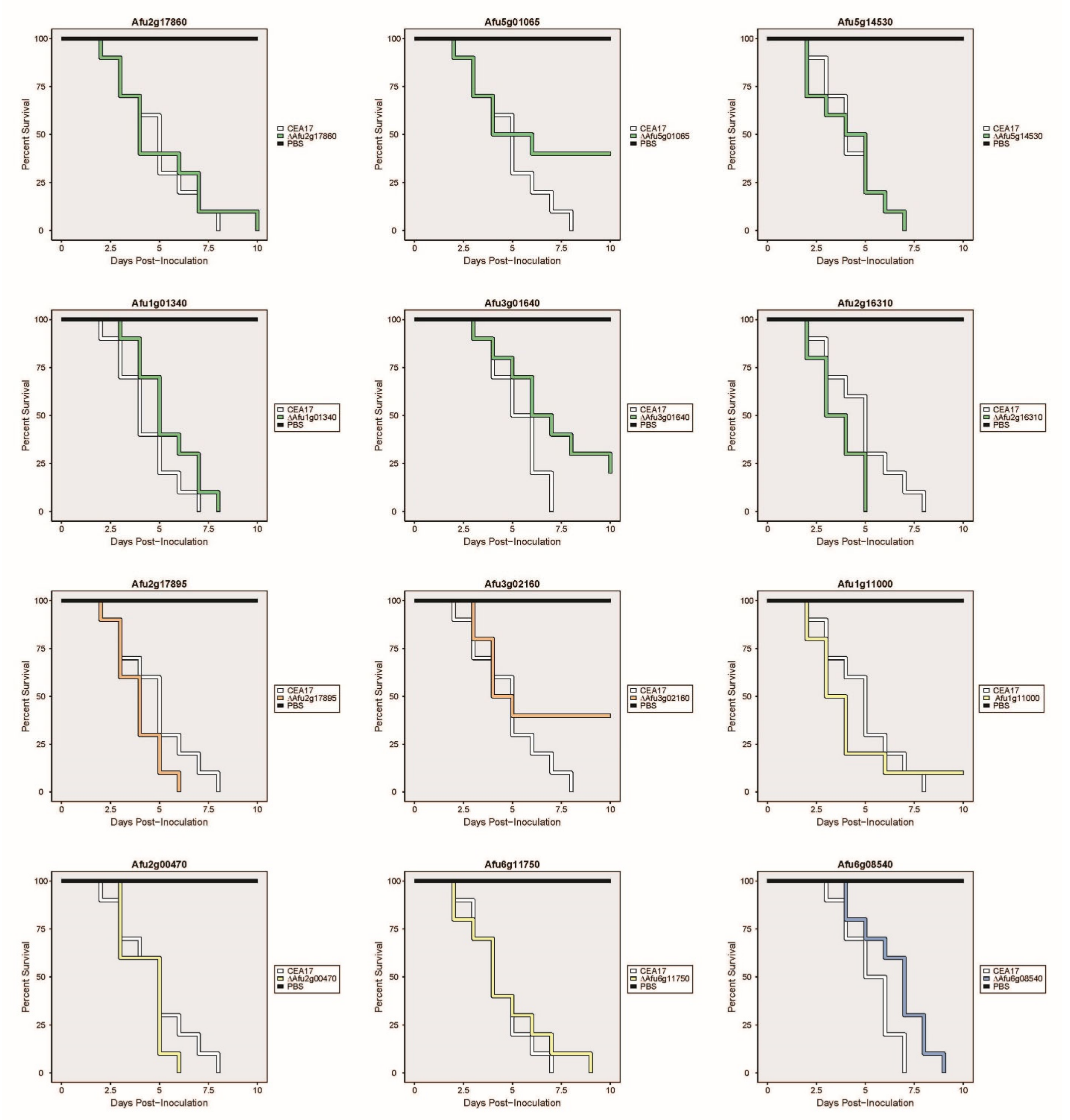


**Supplementary Figure S5. Multiple transcription factors whose evolution differs with respect to pathogenicity do not affect virulence in the greater wax model of disease.** Cumulative survival of *Galleria mellonella* larvae inoculated with phosphate buffered saline (Black), asexual spores (conidia) of the parental strain CEA17 (white), and asexual spores from null mutants of transcription factors with genomic traits related to virulence (colored). Ten larvae were used per inoculation in all assays. Color scheme is the same as in Figure 4. All mutant survival curves shown here were not statistically different ($p>0.05$ in a Log-Rank test) from the CEA17 survival curve.

**Supplementary Tables**

**Table S1. Summary of genomes used in the study.**

**Table S2. Branch information for Figure 1.** Gene gains, losses, and the net gene gain or loss of the branches labeled in Figure 1.

**Table S3. Virulence-related genes shown in Figure 2A.** Note that the order in this table is the same from left to right as in Figure 2A.

**Table S4. Genes that exhibited pathogenicity-related traits.** Gene identifiers (AFUA_#G#####) are for the *A. fumigatu*s strain AF293 member of the gene family except where the *A. fischeri* gene identifier is given (NFIA_######).
